# Gene loss dynamics and T3PKS conservation shape the evolution and pathogenicity of *Rosellinia necatrix*

**DOI:** 10.64898/2026.01.26.701723

**Authors:** Gabriel Quintanilha-Peixoto, Ana Luiza Martins Karl, Dayana K. Turquetti-Moraes, Felipe Rimes-Casais, Aristóteles Góes-Neto, Thiago M. Venancio

## Abstract

Fungal pathogens exhibit remarkable genome plasticity, driven by polyploidy, genome duplication, transposable elements, and niche adaptation. Gene losses often occur in dispensable regions, including in the remarkably dynamic secondary metabolite gene clusters (SMGCs). Within the diverse family Xylariaceae, comprising endophytes, saprotrophs, and phytopathogens, the broad-spectrum pathogen *Rosellinia necatrix* is of major concern, causing white root rot in numerous crops worldwide. Its strategy involves the root infection of weakened plants, tissue colonization, and saprotrophic survival in soil; yet, the genetic basis of this versatility remains poorly understood. Herein, we applied comparative genomics across Xylariaceae to investigate the molecular determinants of *R. necatrix* pathogenicity. We uncovered widespread gene losses in *R. necatrix*, particularly in SMGCs, candidate effectors, and transporter families (MFS and ABC transporters), suggesting a streamlining of its metabolic repertoire during adaptation to diverse hosts. We also identified two highly conserved type III polyketide synthases (T3PKS) across the family, predicted to encode chalcone synthases. Structural modeling and docking analyses support their role in chalcone-related biosynthesis, pointing to an unexpected link between fungal metabolism and plant-associated compounds. Variation in SMGC and carbohydrate-active enzyme (CAZy) repertoires across Xylariaceae further suggests a hemibiotrophic potential for *R. necatrix*, reconciling its capacity for both latent colonization and aggressive necrosis. Our findings establish niche specificity as a key driver of genome reduction in *R. necatrix* and reveal conserved metabolic innovations across Xylariaceae. By integrating gene loss dynamics with secondary metabolism, this work provides new insights into fungal adaptation and pathogenicity, with implications for disease management in perennial and annual crops.

## INTRODUCTION

Fungi exhibit remarkable genome plasticity, with variations in gene content driven by multiple factors. These include changes in ploidy (Albertin & Marullo, 2012), segmental or whole-genome duplications (Van Westerhoven et al., 2024), and the action of transposable elements (Gupta et al., 2023), occasionally enabling adaptation to new hosts or niches (Sharma et al., 2014). In general, gene losses are preferentially confined to dispensable genes, particularly those involved in secondary metabolism (Liang et al., 2018).

The family Xylariaceae is highly diverse, encompassing endophytes, phytopathogens, wood-decay fungi, and other saprophytes (Franco et al., 2022; Samarakoon et al., 2022; U’Ren et al., 2016). Its members occupy a wide array of substrates, ranging from plant hosts to lichen thalli and non-specific wood sources. Within this family, several genera harbor important plant pathogens, including *Biscogniauxia*, *Ustulina*, and *Nemania* (Franco et al., 2022). Notably, Xylariaceae also includes *Rosellinia necatrix*, the causal agent of white root rot in numerous crops, which leads to considerable yield losses in woody, shrub-like, and annual hosts. Documented susceptible species include avocado (Hartley, Engelbrecht & Van den Berg, 2022; Wingfield et al., 2022), rose (Chavarro-Carrero et al., 2024), pear (Shimizu, Kanematsu & Yaegashi, 2018), and soybean (Henning et al., 2014).

Plant pathogens can be classified according to their lifestyle strategies during host interaction. Biotrophic pathogens sustain host vitality while extracting nutrients from the host’s tissues. To preserve host cell integrity, these organisms rely on specialized feeding structures known as haustoria, which are also essential for delivering effector proteins that modulate plant physiology and suppress immune responses (Fei & Liu, 2023). In contrast, necrotrophic pathogens destroy host cells to obtain nutrients, either by penetrating and rupturing cell membranes with invasive hyphae or by inducing programmed cell death while colonizing the apoplast (Mengiste, 2012; Quintanilha-Peixoto et al., 2022). Hemibiotrophic pathogens combine both strategies, initiating infection through a biotrophic phase before transitioning to a necrotrophic stage that completes their life cycle (Plaumann et al., 2018). Beyond visible disease symptoms, these trophic strategies can often be inferred from characteristic genomic features (Hane et al., 2020).

Interestingly, some endophytic fungi may shift toward a pathogenic lifestyle under abiotic stress conditions. This appears to be the case for *Biscogniauxia*, an endophytic genus commonly associated with oak trees (*Quercus* spp.) (Purbaya et al., 2023), which can adopt a necrotrophic phase under elevated temperatures (Costa et al., 2022; Tropf et al., 2025). Similarly, previous studies have reported a saprotrophic-to-necrotrophic lifestyle in *Ustulina deusta* (syn. *Kretzschmaria deusta*), a wood-decaying species (Zmitrovich; Shishlyannikova, 2025).

The pathogenic strategy of *R. necatrix* is characteristic of a broad-range necrotroph. The fungus infects weakened plants through the roots and, once the rot reaches the vascular tissues, ultimately causes host death (Ten Hoopen & Krauss, 2006). Its necrotrophic potential is further supported by its ability to persist in the soil as a saprotroph in the absence of a plant host (Ikeda, Nakamura & Matsumoto, 2005). At the molecular level, necrotrophs heavily rely on effectors, secondary metabolites, and cell wall-degrading enzymes to induce hypersensitivity responses in plants, a defense mechanism that triggers programmed cell death (Rodriguez-Moreno et al., 2018). Following tissue death, necrotrophic pathogens secrete enzymes that degrade host nutrients externally (Zhao et al., 2013).

In this study, we employed comparative genomics to investigate phylogenetic relationships within Xylariaceae, with a particular focus on identifying molecular signatures in *R. necatrix* that may underlie its adaptation to agricultural environments. Our results reveal extensive gene loss in *R. necatrix*, particularly in genes associated with effector activity and secondary metabolite biosynthesis. We identified two highly conserved type III polyketide synthases (T3PKS), likely involved in the synthesis of chalcone-related compounds. Furthermore, the variation in secondary metabolite gene clusters (SMGCs) and carbohydrate-active enzyme (CAZy) repertoires among *R. necatrix* genomes supports the hypothesis of a hemibiotrophic lifestyle.

## METHODS

### Genomic data and quality filtering

A total of 133 genomes from the family *Xylariaceae* were initially analyzed (Table S1). Genome assemblies were retrieved from GenBank and subjected to quality filtering based on BUSCO v5.4.2 completeness (Simão et al., 2015) and standard assembly metrics (including the number of contigs and N50). BUSCO analyses were performed with the Sordariomycetes dataset, which contains 3,817 marker genes. Genomes with less than 70% completeness were excluded from downstream analyses.

### BUSCO-assisted phylogenomic analysis, redundancy filtering, and genome annotation

For an early phylogenetic reconstruction and genome curation, the single-copy orthologs (SCOs) dataset detected by BUSCO using the Sordariomycetes dataset was clustered using OrthoFinder v2.5.5 (Emms; Kelly, 2019) using their default MAFFT module. The phylogenetic reconstruction was conducted with IQTREE2 v2.1.4-beta (Minh et al., 2020) under the “TEST” mode for best-fit model selection. Genomes lacking clear genus-level classification or those positioned outside the *Nemania/Rosellinia* clade were removed from further analysis.

The remaining genomes were re-annotated with the Funannotate pipeline v1.8.15 (Li; Wang, 2021). Transcript and protein evidence databases were generated from publicly available mRNA and protein sequences available on GenBank (accessed in January 2024). Redundancy was removed using CD-HIT v4.8.1 (Fu et al., 2012) with 100% identity, and the resulting non-redundant datasets were supplied to Funannotate. Genomes were also annotated for the presence of SMGCs with antismash v7.1.0 (Blin et al., 2021). Functional annotation of genes of interest was performed using InterProScan (Jones et al., 2014) and dbCAN3 (Zheng et al., 2023). Species trophic modes were inferred with Catastrophy (Hane et al., 2020).

### Phylogenomics, gene repertoire, and gene loss

Gene repertoire visualization was performed using the ortholog family abundance matrix generated by OrthoFinder, implemented through in-house R scripts. A second phylogenetic reconstruction was obtained, this time using SCOs obtained from the reannotated genomes, instead of a BUSCO marker gene dataset. In this analysis, multiple sequence alignments obtained with OrthoFinder were used to reconstruct a phylogenetic tree with IQTREE2, as described above. Gene family expansions and contractions were assessed with CAFE v5 (Mendes et al., 2021), and statistically significant changes (p-value < 0.05) were filtered using in-house bash scripts. Ortholog families were classified into gene categories based on their frequency across genomes: core (present in all genomes), softcore (>95%), shell (2–95%), and cloud (<2%), following Matthews et al. (2024).

### Protein structure modelling, docking, and virtual screening

Target protein sequences were searched against the RefSeq database using BLASTp (online available version) (Camacho et al., 2009). Tertiary structure predictions were generated using I-TASSER (Yang et al., 2015) and AlphaFold (Jumper et al., 2021). Model quality was assessed using MolProbity (Williams et al., 2018), and structure-based function prediction was performed with the DALI server (Holm et al., 2023).

Protein structures selected for docking were prepared for molecular modeling using the PrepWizard tool within the Maestro suite (Schrödinger Release 2025-2: Protein Preparation Workflow; Epik, Schrödinger, LLC, New York, NY, 2024; Impact, Schrödinger, LLC, New York, NY; Prime, Schrödinger, LLC, New York, NY, 2025). Preparation included adding missed atoms, adding hydrogen atoms, optimizing protonation states at pH 7.0, and energy minimization with the OPLS4 force field to resolve structural inconsistencies.

A customized ligand library was constructed by combining Sordariomycetes secondary metabolites obtained from FungiDB (Basenko et al., 2018) with stillbenes, chalcones, and hydroxymethylglutaryl-CoA (HMG-CoA) precursors and derivatives from ChEBI (Hastings et al., 2016). All ligands were standardized with the LigPrep module (Schrödinger Release 2025-2: LigPrep, Schrödinger, LLC, New York, NY, 2025; Johnston et al., 2023), including adjustment of protonation states, stereochemistry, and tautomeric forms.

Protein models and ligand libraries were imported into Maestro for molecular docking and virtual screening analyses. All computational procedures were conducted using standardized parameters to ensure reproducibility. Binding cavities were initially explored with CB-Dock2 (Liu et al., 2022). Blind docking of chalcone (CHEBI:27618) was performed using AutoDockVina (Eberhardt et al., 2021). We selected the two best-ranked cavities for virtual screening experiments.

Virtual screening experiments were conducted with Glide (Schrödinger Release 2025-2: Glide, Schrödinger, LLC, New York, NY, 2025), targeting the identified binding cavities. Ligands were screened using the High-Throughput Virtual Screening (HTVS) protocol, and the best docking poses were selected based on Glide scores and visual inspection in PyMOL (The PyMOL Molecular Graphics System, Version 3.0, Schrödinger, LLC.).

## RESULTS

### Xylariaceae phylogenomics and gene repertoire

We conducted a comparative genomic analysis of 44 high-quality Xylariaceae genomes (Figure 1A, S1). BUSCO completeness (Simão et al., 2015), assessed using the Sordariomycetes database, ranged from 72.2% in *R. necatrix* R10 to 98.3% in *R. necatrix* W97. Most assemblies are organized in fewer than 2,000 contigs, except for *Xylaria arbuscula* VT107, which has 3,895 contigs. The proportion of repetitive (masked) regions varied substantially, from 5.25% in *Xylaria arbuscula* VT107 to 33.6% in *Poronia punctata* CBS 180.79. After reannotation, the number of protein-coding genes varied from 9,513 in *P. punctata* CBS 180.79 to 13,901 in *Nemania abortiva* FL1152. All reannotation files are available at https://github.com/gqpx/xylariaceae.

**Figure 1:**
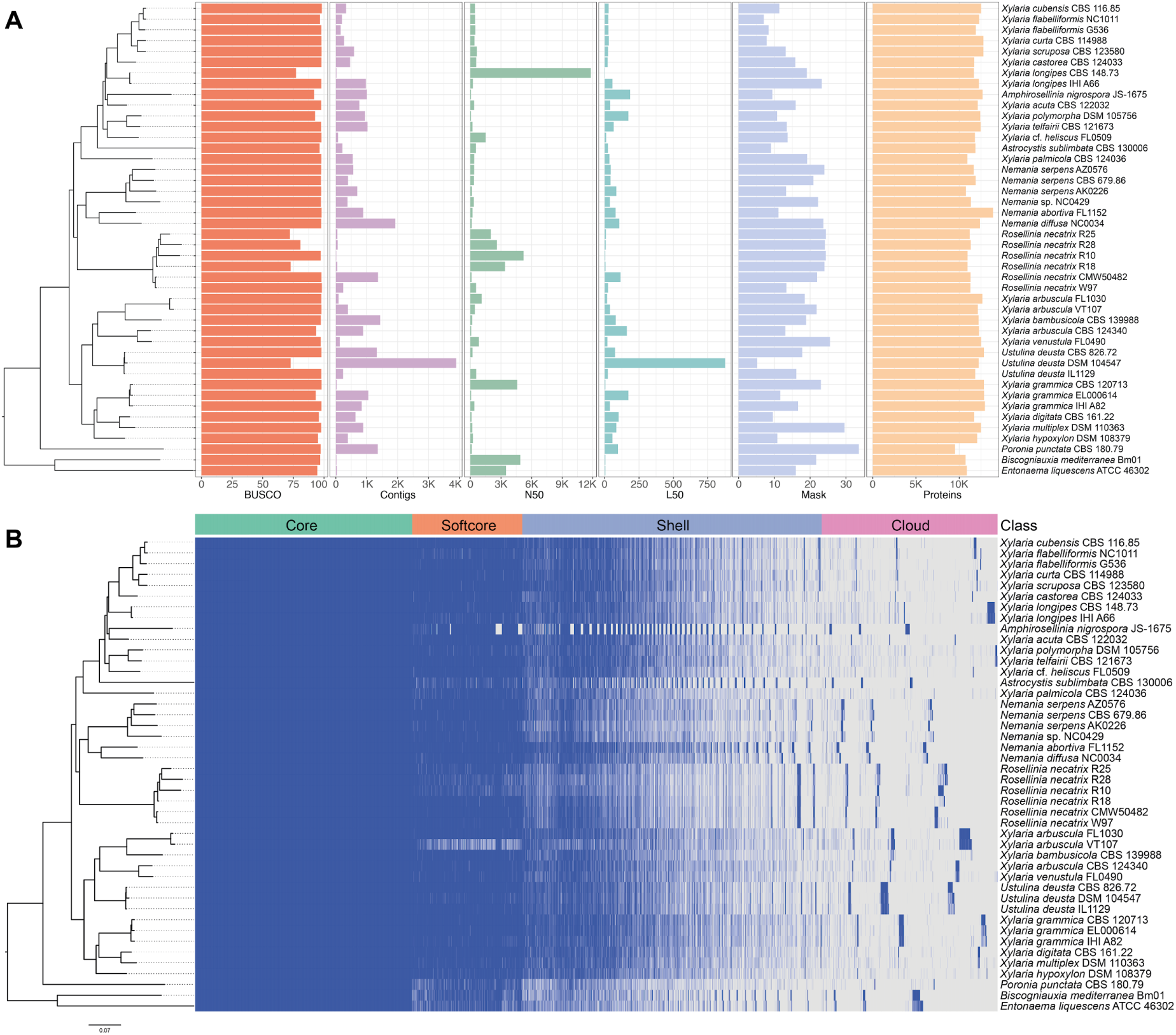
Phylogenomics of Xylariaceae. **A:** Phylogenomic tree with genome assembly and annotation metrics, including BUSCO completeness, number of contigs, N50, L50, percentage of masked/repetitive regions, and the number of proteins. **B:** Phylogenomic tree with gene distribution across genomes, showing the proportion of core, softcore, shell, and cloud gene families.

Our gene repertoire analysis, performed with OrthoFinder (Emms; Kelly, 2019), identified 4,705 core gene families (present in all genomes), 2,380 softcore families (≥95% of genomes), 6,033 shell families (found in 6 to 41 genomes), and 4,260 cloud families (present in 1 to 5 genomes) (Figure 1B). Gene repertoire proportions were adapted from Matthews et al. (2024). A detailed annotation of the *R. necatrix* cloud genome is presented in Figure S2.

We used OrthoFinder SCOs to perform a phylogenomic reconstruction and infer a species tree with IQ-TREE2 v2.1.4-beta (Minh et al., 2020). The analysis, based on 1,425 SCOs and employing the Q.mammal+F+I+G4 substitution model (best-fit model selected by IQ-TREE), recovered monophyletic clusters for all formal genera, except *Xylaria*, a genus traditionally used to describe wood-decay ascomycete fungi (Stadler et al., 2013).

### Trophic mode of Rosellinia necatrix

The trophic mode of all genomes was obtained with Catastrophy (Hane et al., 2020). For *R. necatrix,* the primary trophic mode was predicted as polymertroph (broad or narrow range), which corresponds to necrotrophy. However, most of the genomes were also classified as mesomertrophs (equivalent to hemibiotrophs) with a score greater than 0.9 (Table 1). Therefore, the trophic mode of *R. necatrix* remains inconclusive between necrotrophy and hemibiotrophy. The full annotated CAZy repertoire (used by Catastrophy for trophic mode determination), obtained with dbCAN3 (Zheng et al., 2023), is provided in Table S2.

**Table 1:** Trophic mode scores of *R. necatrix* genomes predicted with Catastrophy (Hane et al., 2020).

| Isolate | PB | PN | MeE | MeI | Mo1 | Mo2 | Me3 | S | V |
| --- | --- | --- | --- | --- | --- | --- | --- | --- | --- |
| CMW50482 | 0.812 | 1 | 0.688 | 0.947 | 0.380 | 0 | 0.348 | 0.511 | 0.801 |
| R10 | 1 | 0.826 | 0.908 | 0.673 | 0.601 | 0 | 0.573 | 0.751 | 0.446 |
| R18 | 0.910 | 1 | 0.768 | 0.957 | 0.467 | 0 | 0.421 | 0.604 | 0.792 |
| R25 | 1 | 0.977 | 0.840 | 0.906 | 0.536 | 0 | 0.490 | 0.685 | 0.682 |
| R28 | 1 | 0.930 | 0.900 | 0.813 | 0.581 | 0 | 0.553 | 0.728 | 0.579 |
| W97 | 0.822 | 1 | 0.686 | 0.948 | 0.385 | 0 | 0.348 | 0.516 | 0.793 |

Abbreviations: PB, polymertroph (broad spectrum); PN, polymertroph (narrow spectrum); MeE, extracellular mesotroph; MeI, intracellular mesotroph; Mo1, monomertroph type 1; Mo2, monomertroph type 2; Mo3, monomertroph type 3; S, saprotroph; V, vascularotroph.

### Ortholog family expansion and contraction

Using CAFE v5 (Mendes et al., 2021), we identified multiple ortholog family expansions and contractions across Xylariaceae. A striking pattern emerged in the clade comprising all *R. necatrix* assemblies, in which a total of 1,028 gene family contractions were detected. Of these, 795 were statistically significant (p < 0.05), with 778 (97.9%) corresponding to complete gene family losses across *R. necatrix* genomes and 17 reflecting partial losses restricted to a subset of assemblies. In contrast, expansions were more than six times less frequent, with 141 cases in *R. necatrix*, 120 of which were statistically significant (Figure 2, S3).

**Figure 2:**
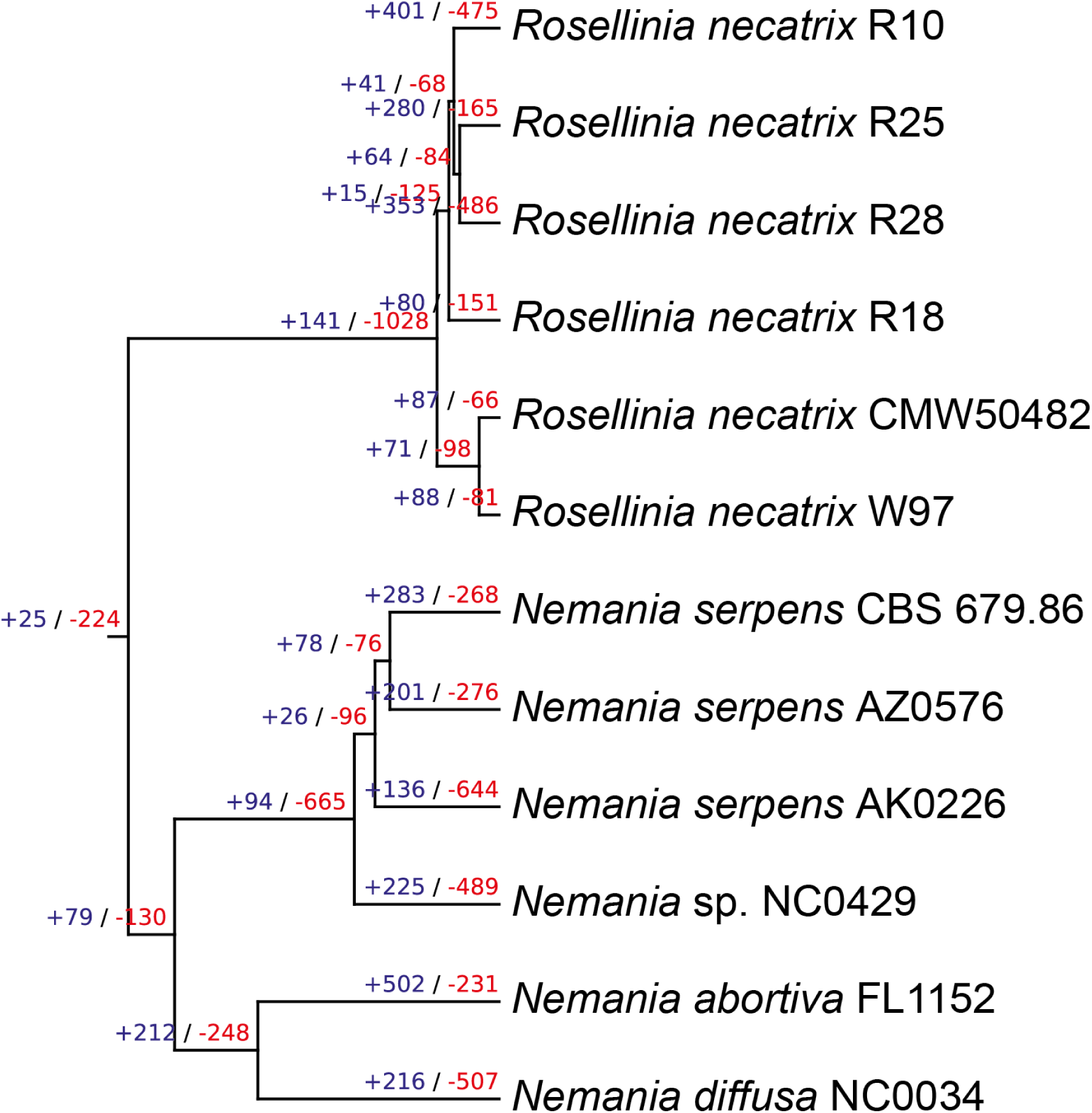
Ortholog family expansions and contractions inferred with CAFE v5. Blue and red values indicate gene family gains and losses, respectively.

### Tracking gene losses in *R. necatrix*

Our ortholog analysis with CAFE v5 identified 778 orthologous families absent from all *R. necatrix* genomes. Among these, 110 families contain orthologs present in all *Nemania* genomes, suggesting that their loss predated the diversification of *R. necatrix* strains. To minimize biases associated with genome fragmentation, we selected *Nemania abortiva* FL1152 as a reference for describing the ortholog gene families absent in *R. necatrix*.

Among the 768 families absent in *R. necatrix*, 551 are retained in *N. abortiva*, comprising a total of 768 genes. Amongst the 110 families found across all *Nemania* genomes, 64 could be functionally annotated with InterProScan analysis of their *N. abortiva* orthologs (Table 2). Notably, 158 of these *N. abortiva* orthologs (20%) are located within SMGCs, as annotated with antiSMASH; however, recognizable protein domains or families could not be detected in most of these. SMGC-related functions were recovered through functional annotation (Table 2). Beyond this group, we highlight the second most frequent functional category among the genes absent in *R. necatrix*: transmembrane transporters (Table 2, Figure 4), particularly ABC transporters. Six other interesting regions of genomic losses were analyzed, showing a similar pattern (Figure S4).

**Table 2:**
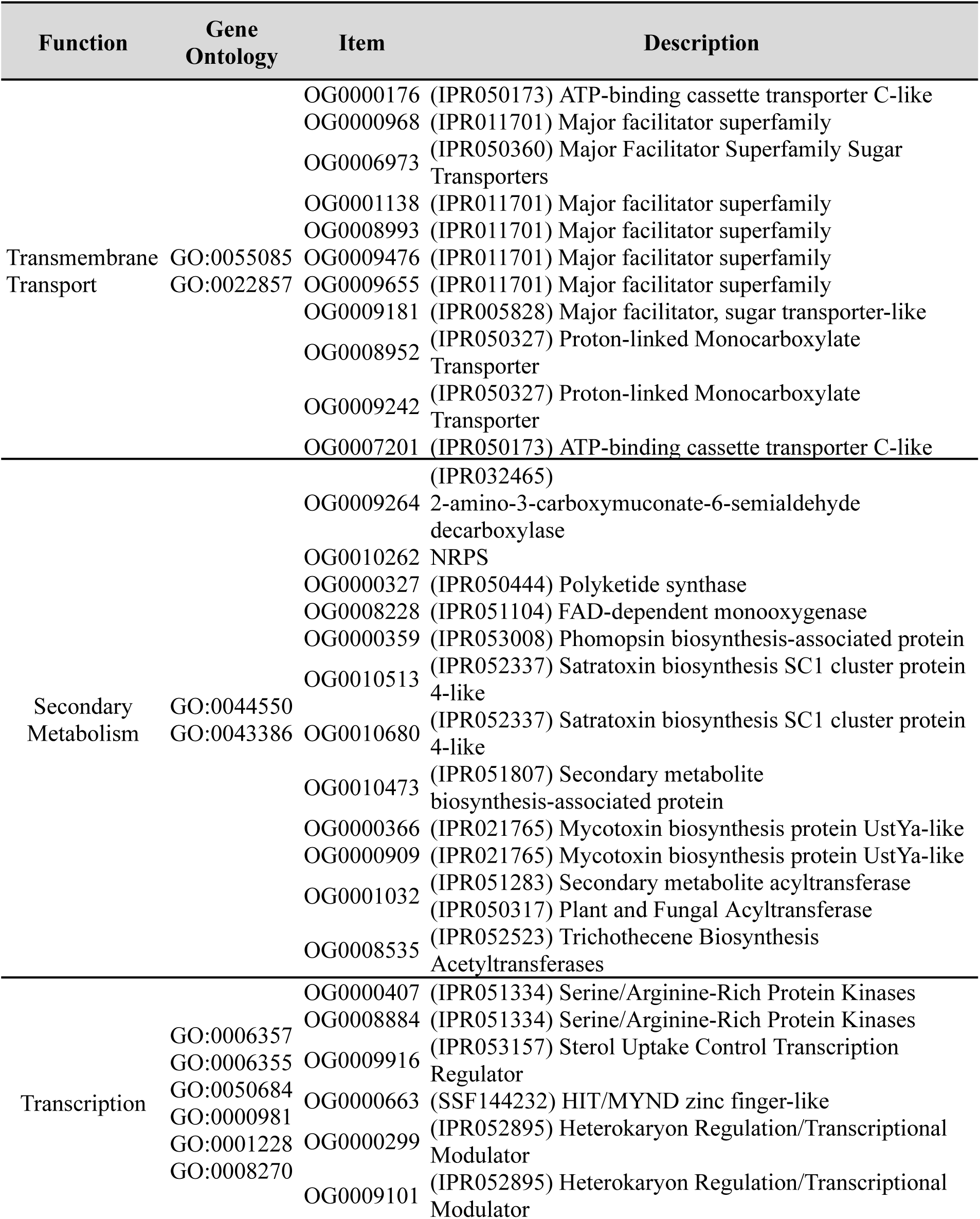
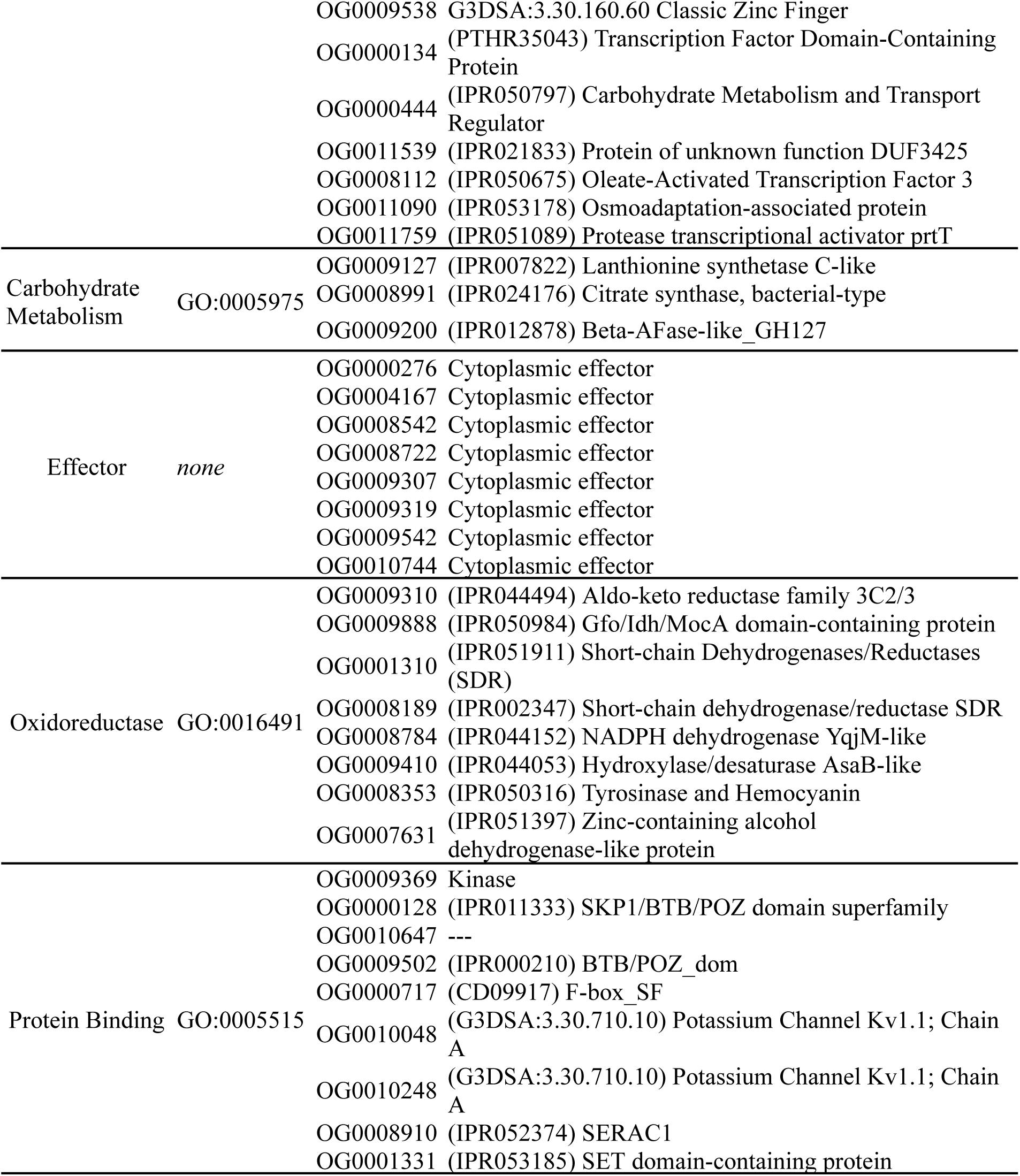
***N. abortiva* FL1152 orthologs corresponding to gene families lost in *R. necatrix*.**

### Type III polyketide synthase conservation in Xylariaceae secondary metabolite gene clusters

Analysis of SMGCs in Xylariaceae genomes annotated with antismash v7.1.0 (Blin et al., 2021) revealed a conserved pattern of T3PKS clusters. All species analyzed contain two T3PKS clusters, except *Biscogniauxia mediterranea*, *Entonaema liquescens*, and one strain of *Xylaria longipes*. A sequence-similarity network based on the main annotated biosynthetic gene resolved two distinct protein groups. Cluster I comprises chalcone synthases ranging from 423 to 480 amino acids in length (mean: 475 aa), except the *Amphirosellinia nigrospora* protein, which is truncated at 209 aa. Cluster II corresponds to chalcone/stilbene synthases ranging from 345 to 395 aa in length (mean: 384 aa), again with a shorter *A. nigrospora* protein of 247 aa. Notably, *B. mediterranea* and *E. liquescens* each harbor only a single protein in Cluster I, while *X. longipes* has entirely lost its Cluster I gene (Figure 3). The complete SMGC annotation for Xylariaceae is provided in Table S3.

**Figure 3:**
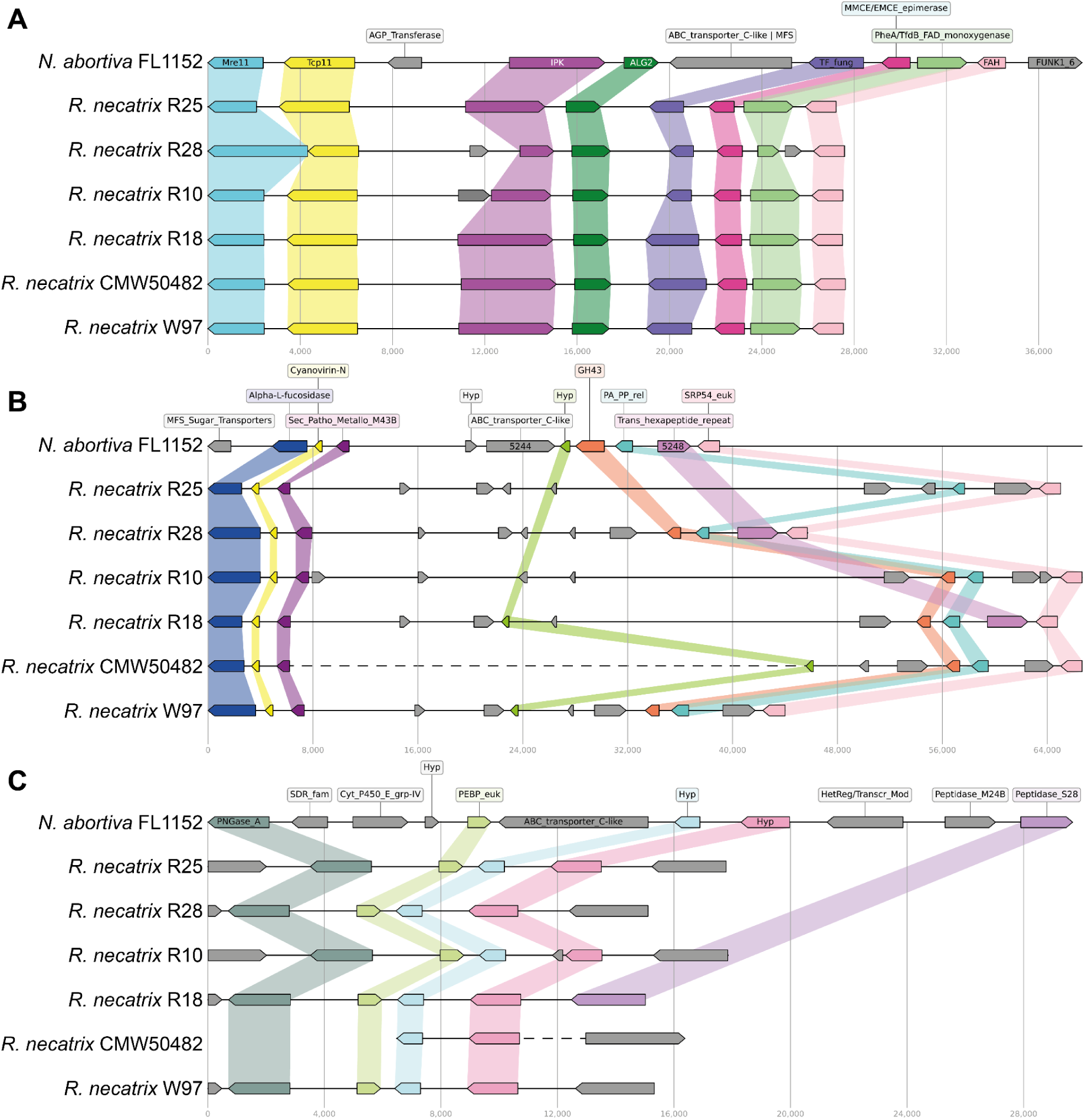
Gene losses in *R. necatrix* genomes compared with *N. abortiva* FL1152. Each arrow represents a single gene; colors and connecting structures indicate similarity, while genes in grey displayed no detectable similarity between *R. necatrix* and *N. abortiva*. Gene labels are shown for the reference genome. Dashed lines indicate contig gaps. **A:** OG0000176 (main gene: *nabor_001615*). **B:** OG0000176 (main gene: *nabor_005244*) **C:** OG0007201 (main gene: *nabor_012118*).

**Figure 4:**
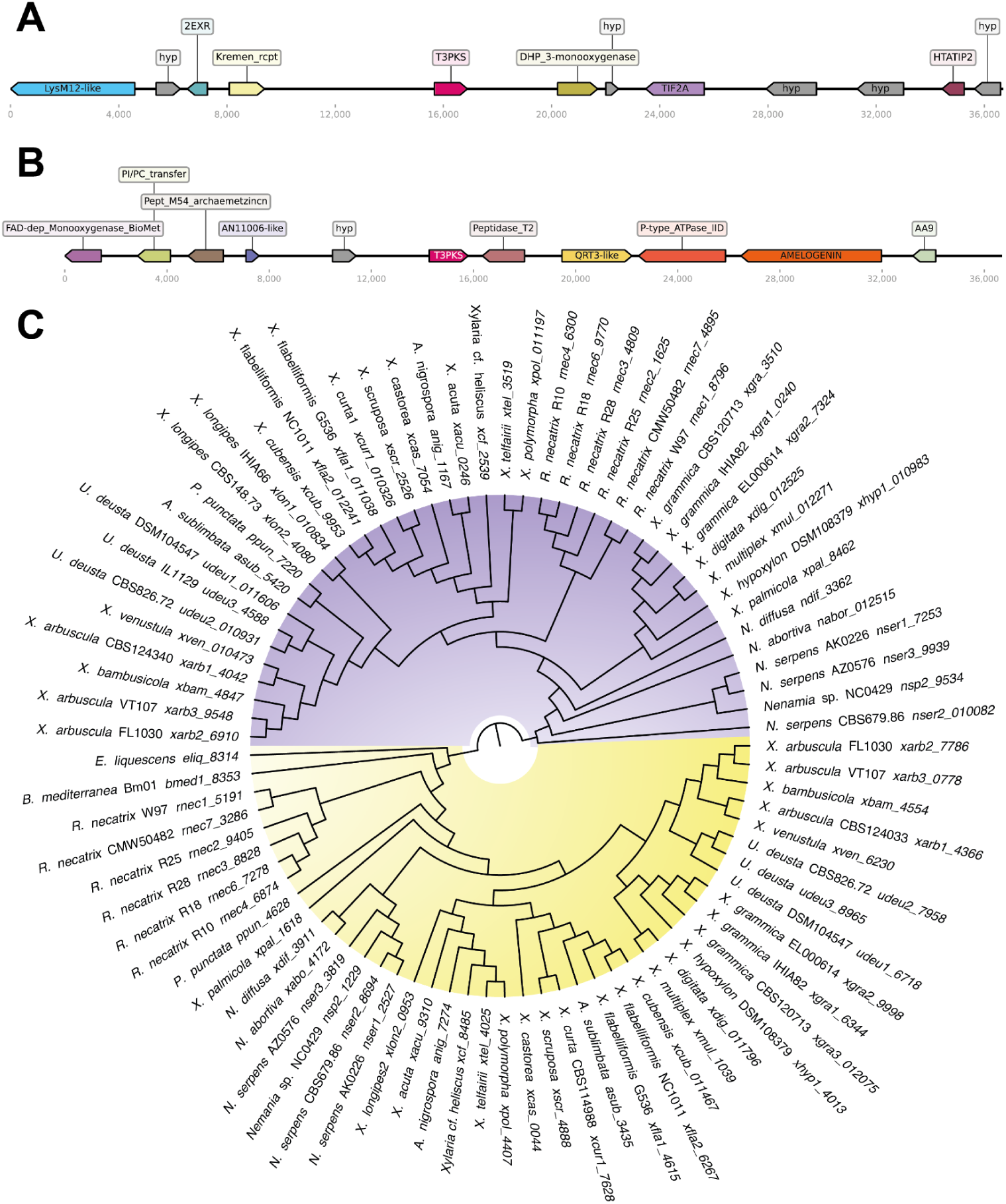
Distribution of secondary metabolite gene clusters (SMGC) in Xylariaceae. **A:** Genomic context of T3PKS Cluster I in *Rosellinia necatrix* CMW50482. **B:** Genomic context of T3PKS Cluster II in *R. necatrix* CMW50482. **C:** Phylogenetic tree of type III polyketide synthases (T3PKS).

### Modelling chalcone synthases in *R. necatrix*

A BLASTp search against the nr database (using default values) (Camacho et al., 2009) revealed that *rnec7_4895* and *rnec7_3286* (Figure 4C) share sequence similarity with two T3PKS (WP_168884327.1 and WP_235288137.1, respectively). InterProScan annotation (Jones et al., 2014) further indicated that, in addition to T3PKS, these enzymes were annotated as HMG-CoA synthases (PTHR11877). For *rnec7_3286*, three predicted malonyl-CoA-binding regions were identified at residues 210, 269, and 316-318 (CD00831: CHS_like, encompassing chalcone and stilbene synthases, plant-specific polyketide synthases, and related enzymes). Both predicted proteins also contain conserved N-terminal (IPR016039) and C-terminal (IPR016039) domains of chalcone synthases.

To investigate their structural features, three-dimensional models were generated using I-TASSER (Yang et al., 2015) and AlphaFold (Jumper et al., 2021). Model refinement was guided by Ramachandran plot evaluations with MolProbity (Williams et al., 2018) and, subsequently, optimized with Maestro. Two main cavities were predicted for each protein (Table S4-S5).

The final 3D models (Figure 5A-B) aligned closely with the template structure 3E1H from PDB, forming monomeric α/β proteins consistent with their annotated domains. Structural comparison with the DALI server revealed strong similarity to a previously characterized T3PKS (PDB ID: 3EUQ). Considering that chalcones are polyketide derivatives, this structural alignment corroborates the BLASTp annotations.

**Figure 5:**
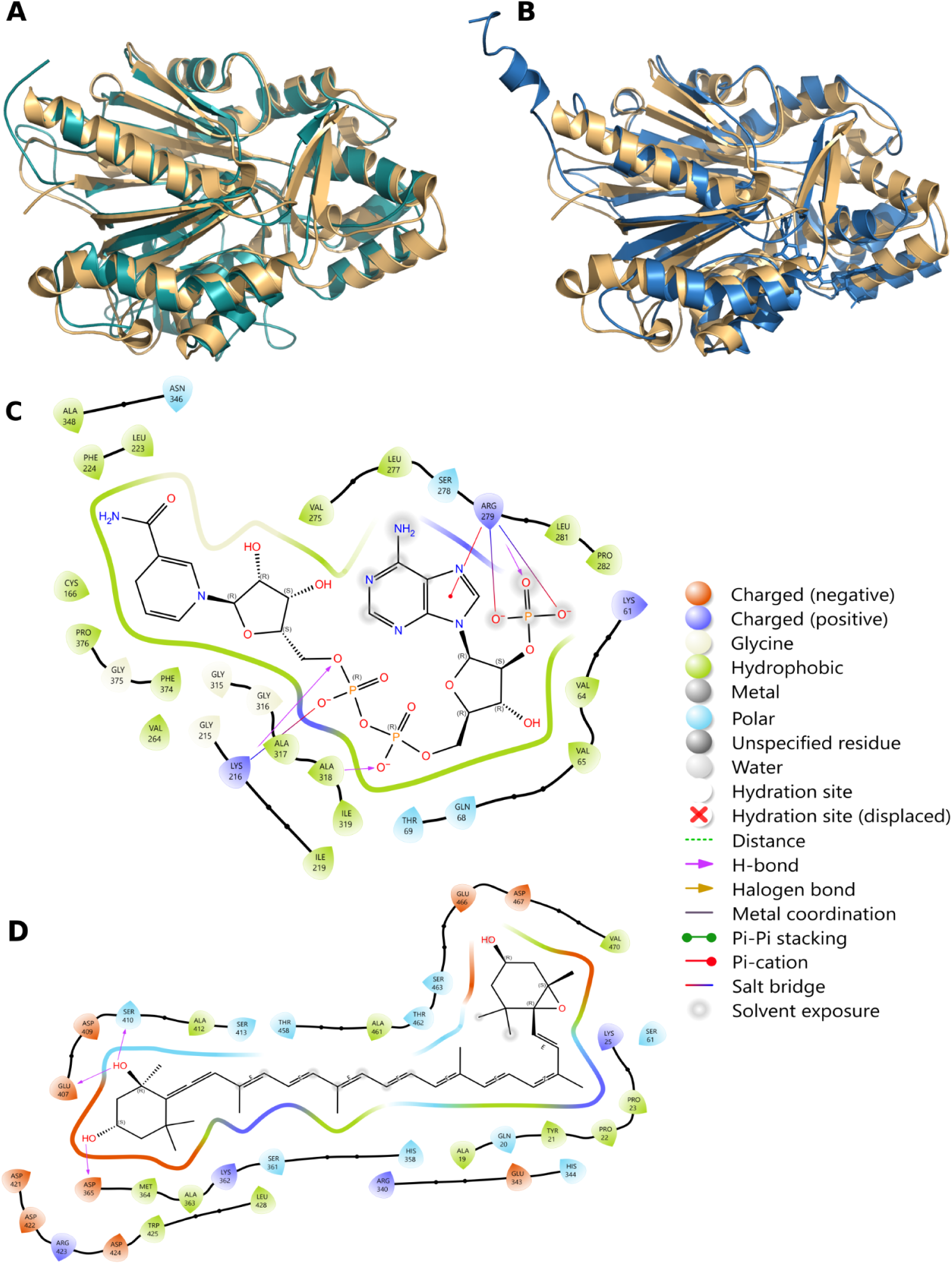
Molecular modeling of chalcone synthases from *Rosellinia necatrix*. **A:** Predicted structure of *rnec7_3286* (cyan) aligned with its template 3E1H (light orange). **B:** Predicted structure of *rnec7_4895* (blue) aligned with the same template 3e1h (light orange). **C:** Predicted binding pose of Coenzyme A within cavity 1 of *rnec7_4895*, with the corresponding Maestro 2D ligand interaction scheme. **D:** Predicted binding pose of malonyl-CoA within cavity 1 of *rnec7_3286*, with the corresponding Maestro 2D ligand interaction scheme.

For functional investigation, we assembled a customized ligand library consisting of 837 compounds from FungiDB associated with Sordariomycetes species, which was supplemented with 183 chalcones, 62 stilbenes, and 7 acetyl-CoA-related compounds from ChEBI, totalling 1,119 unique ligands. Three-dimensional ligand structures were retrieved in SDF format from ChEBI and prepared with LigPrep, yielding a final library of 6,203 conformations for subsequent docking analysis.

For *rnec7_3286*, cavity 1 exhibited the most favorable binding energies. The top-ranked compounds from virtual screening (VS) were further filtered based on their reported occurrence either in fungi or chalcone metabolism (Table 3). Notably, malonyl-CoA⁵⁻ ranked among the top three compounds, with a binding energy of −10.971 kcal/mol (Figure 5C). Conversely, *rnec7_4895* showed weaker binding energies compared to *rnec7_3286*, with scores for its main cavity consistently above −8 kcal/mol (Table 3). Coenzyme A was also among the top-ranked compounds in the *rnec7_3286* VS experiments (binding energy: −7.862 kcal/mol; Figure 5D), while malonyl-CoA ranked within the top 11 compounds (−7.407 kcal/mol). The binding residues predicted for malonyl-CoA binding in the InterProScan annotation did not correspond to any relevant cavity in our modelling analysis (Figure S5).

**Table 3:**
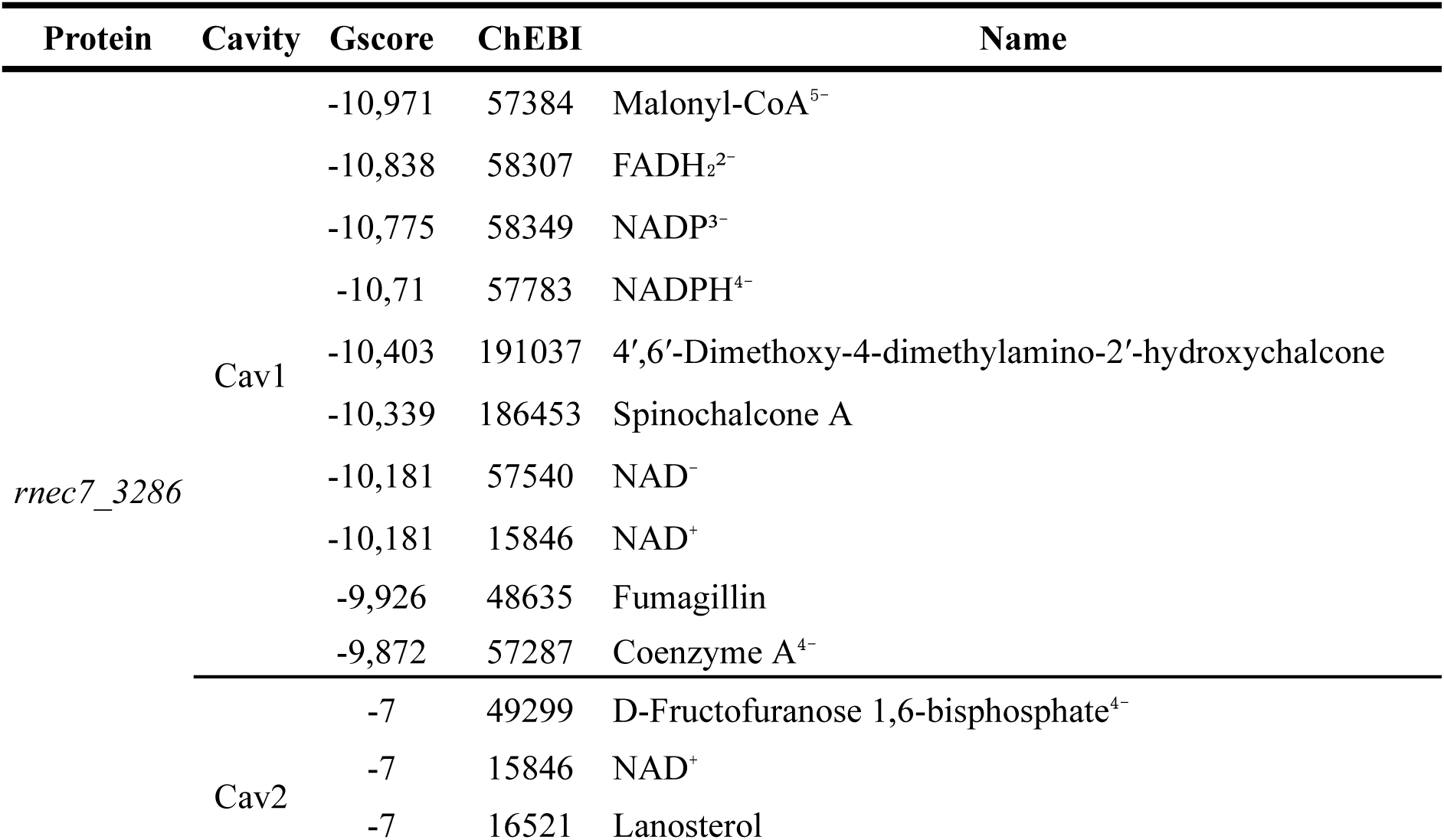
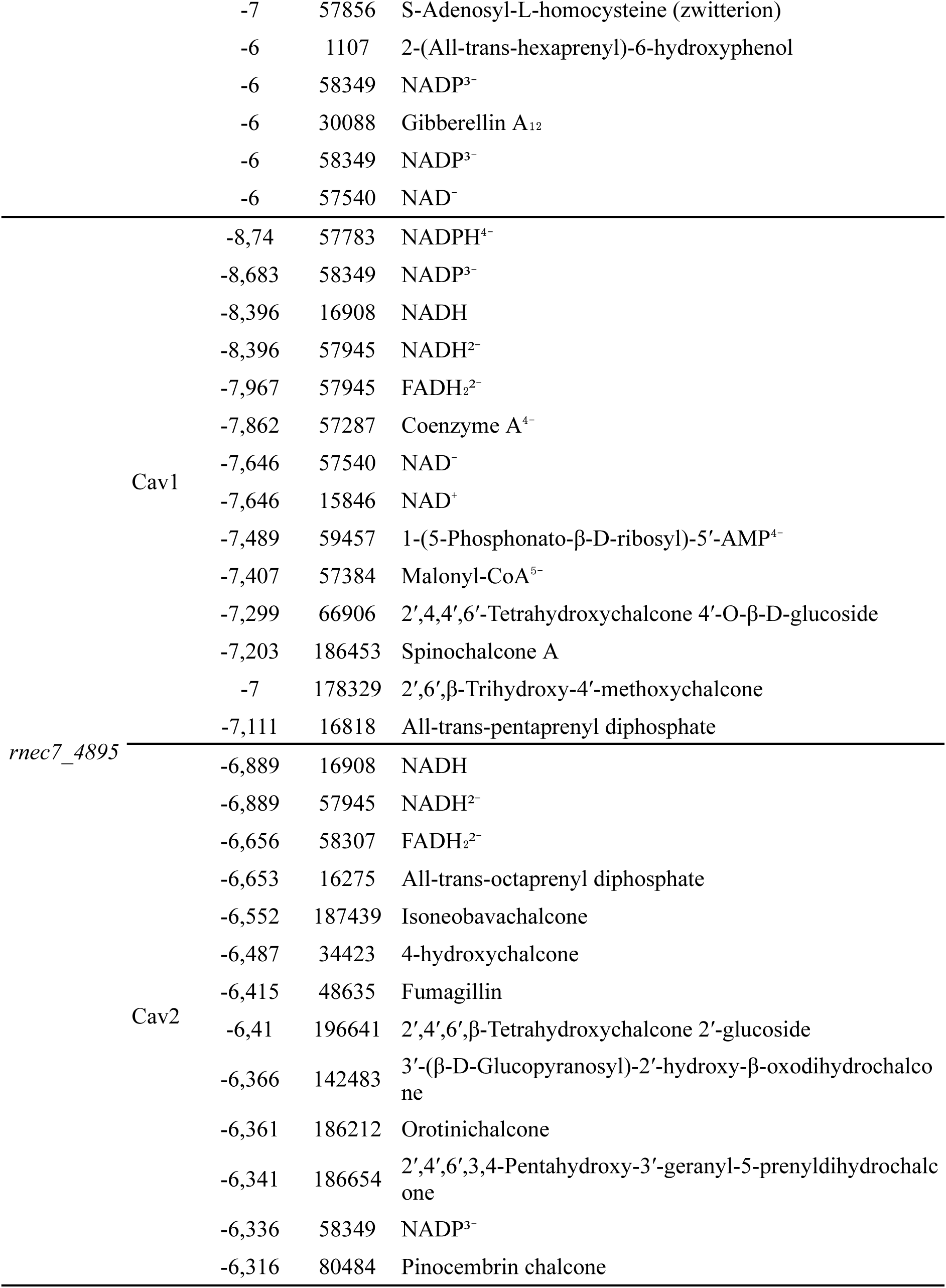
Top-ranked compounds identified through virtual screening in the two main cavities of *Rosellinia necatrix* chalcone synthases *rnec7_3286* and *rnec7_4895*.

In addition to these results, several other compounds were successfully docked to both proteins. Twelve chalcone-derived molecules ranked among the top ligands for the main cavity of *rnec7_3286* and both cavities of *rnec7_4895*. Only two stilbenes were represented among the top-ranked compounds for *rnec7_3286*. In contrast, HMG-CoA, acetyl-CoA, acteoacetyl-CoA, and related compounds did not achieve high-ranking scores (Table S6). Detailed results of cavity scoring and VS outcomes are provided in Table 3 and Table S6.

## DISCUSSION

*R. necatrix* is an important necrotrophic fungal pathogen belonging to the family Xylariaceae (Chavarro-Carrero et al., 2024), and it is the causative agent of root rot in a broad range of plant hosts (Hartley, Engelbrecht, & Van den Berg, 2022; Shimizu, Kanematsu, & Yaegashi, 2018; Wingfield et al., 2022). Our study addresses gaps in the understanding of gene loss and conservation at both the species and family levels. Although we selected a few functional protein groups for in-depth analysis (Table 2-3, Figure 3-5), it is evident that *R. necatrix* genomes contain extensive regions of gene loss compared with their closest relatives, particularly those from the genus *Nemania* (Figure 2), which includes both endophytic and pathogenic species (Apurillo et al., 2024; Demir et al., 2025; Medina et al., 2019). Strikingly, the genes lost in *R. necatrix* are primarily associated with context-dependent functions rather than broad, conserved functions. Among the annotated orthologous families, at least 20 correspond to genes associated with seven distinct molecular functions in fungi, including transmembrane transporters, effector proteins, oxidoreductases, and genes related to secondary metabolism, transcription, carbohydrate metabolism, and protein binding (Table 2). Considering that some of these classes are expressed only in certain contexts, rather than universally, this observation aligns with the findings of Coulombe-Huntington & Xia (2017) with *S. cerevisiae*, which demonstrated that such lost and gained genes are often subject to complex transcriptional regulation.

SMGCs generally display narrow taxonomic distributions (Rokas, Wisecaver & Lind, 2018). Nevertheless, the orthogroups analyzed here are broadly conserved across Xylariaceae (particularly in *Nemania*), yet consistently absent in *R. necatrix*. This pattern supports the previous hypothesis that SMGC gene losses contribute to fungal adaptation to plant hosts (Rokas, Wisecaver, & Lind, 2018; Slot, 2017; Wisecaver & Rokas, 2015). Nonetheless, this raises an unresolved question for *R. necatrix*, given its classification as a broad-range necrotroph. The genomes included in our study were isolated from diverse hosts, including rose in Mexico (Chavarro-Carrero et al., 2024), *Pyrus pyrifolia,* and *Persea americana* (Wingfield et al., 2022; Shimizu, Kanematsu, & Yaegashi, 2018).

Interestingly, some families lost in *R. necatrix* correspond to gene deletions reported in *Colletotrichum* (Liang et al., 2018), which include transcriptional regulators, transporters, and genes related to secondary metabolism. As well as *R. necatrix*, *Colletotrichum* species are broad-spectrum plant pathogens. A key distinction, however, lies in its trophic strategy; whereas *Colletotrichum* species are established hemibiotrophs, our results suggest that *R. necatrix* may display either necrotrophic or hemibiotrophic modes (Table 1). Trophic strategy is a fundamental determinant of infection and colonization processes in phytopathogens and is closely tied to their genomic content (Zhao et al., 2013). Indeed, gene loss has been documented in several other phytopathogens, including *Melanopsichium pennsylvanicum* (Sharma et al., 2014), *Colletotrichum higginsianum*, *Magnaporthe oryzae*, *Sclerotinia sclerotiorum*, and *Blumeria graminis* (Spanu et al., 2010), underscoring the role of gene loss as a driver of niche adaptation.

### *Rosellinia necatrix* lost several transmembrane transporters

Ortholog family expansions and contractions with CAFE v5 (Mendes et al., 2021) revealed extensive gene losses across all *R. necatrix* genomes. Despite the growing availability of genome assemblies and consistent findings in previous studies (Coulombe-Huntington; Xia, 2017; Liang et al., 2018; Sharma et al., 2014; Spanu et al., 2010), gene loss in fungi remains a relatively understudied phenomenon.

In our dataset, some *R. necatrix* isolates displayed below-average genome completeness, although this was not correlated with assembly fragmentation. For instance, isolates R28 and R10 presented 44 and 50 contigs with N50 values of 3.4 and 2.0 Mb, respectively. These metrics are comparable to those of *X. grammica* EL000614 (25 contigs, 4.6 Mb N50) and *X. hypoxylon* CBS 122620 (88 contigs, 3.9 Mb N50). Nonetheless, those *R. necatrix* isolates exhibited only ∼72% BUSCO completeness, compared with 96.6% in *X. hypoxylon* and 98.1% in *X. grammica*. Although such discrepancies are present, our CAFE results were consistent, indicating a total of 779 gene family deletions in the *R. necatrix* clade, regardless of genome assembly quality.

Among the ortholog families absent in *R. necatrix* but present in *N. abortiva*, 11 were identified as transmembrane transporters (Table 2). Those were annotated with Gene Ontology terms GO:0055085 (transmembrane transport) and GO:0022857 (transmembrane transporter activity). The lost transporters fall into five distinct protein families. Four belong to the Major Facilitator Superfamily (MFS), including general MFS transporters (IPR011701), sugar transporters (IPR050360), sugar transporter-like (IPR005828), and proton-linked monocarboxylate transporters (IPR050327). Those proteins function as secondary transporters, using electrochemical gradients rather than ATP to mediate substrate transport (Pozdnyakov et al., 2023; Quintanilha-Peixoto et al., 2022). The fifth family consists of ATP-binding cassette transporter C-like (IPR050173). In fungi, these proteins are classified into subfamilies ABCA to ABCG and ABCI (Víglaš; Olejníková, 2021). The detected ortholog families in *R. necatrix* belong to the ABCC subfamily, also known as Multidrug Resistance Proteins (MRPs). Members of this group typically participate in the efflux of xenobiotics and secondary metabolites, contributing to stress tolerance and pathogenicity. Although this subdivision has been described based on the organization of conserved domains (Kovalchuk & Driessen, 2010), detailed functional characterization of fungal MRPs remains largely elusive.

In *R. necatrix* genomes, the loss of ABCC genes coincided with the contraction of their surrounding genomic regions (Figure 4A, C). In one case, the ABCC gene region found in *N. abortiva* was similar to a longer, gene-sparse region in *R. necatrix* (Figure 4B). Even though the genomic context around the lost ABCC gene was not strictly conserved, neighboring genes were generally syntenic across species, except for a hypothetical protein adjacent to *nabor_005244* in OG0000176 and a peptidase S28 in OG0007201, which displayed differential conservation among *R. necatrix* isolates.

Based on the known functions of MRPs and the broader patterns described by Coulombe-Huntington & Xia (2017) for variable genes in fungi, we hypothesize that the loss of these transporters did not negatively impact *R. necatrix* fitness. This could be explained either by the absence of their target bioactive compounds in the typical host environments of *R. necatrix* or by functional redundancy provided by other genes.

### Two type III polyketide synthases are highly conserved in Xylariaceae

Approximately 40% of the Xylariaceae gene repertoire is highly conserved throughout the family (present in over 95% of the included genomes). Among these conserved ortholog families are two T3PKS, both annotated as HMG-CoA synthases (PTHR11877). T3PKS are relatively uncommon in fungi compared to T1PKS (Shi-Kunne et al., 2019), a pattern also reflected in our dataset (Table S3). Although the genomic context around those genes is not conserved, T3PKS enzymes are known to produce diverse bioactive compounds in fungi, including hypocrellin (Ren et al., 2020), as well as tri-, tetra-, or pentaketide pyrones and resorcinols derived from fatty acyl-CoA substrates, particularly malonyl-CoA (Sokolova et al., 2024).

Molecular modeling and docking of the two HMG-CoA synthases revealed an intriguing duality. Both predicted proteins carry the PTHR11877 family signature, classically associated with the synthesis of HMG-CoA from acetyl-CoA as an intermediate in ergosterol biosynthesis and other metabolic pathways (Sayari et al., 2021). Both proteins retain conserved N- and C-terminal domains typical of chalcone synthases; notably, *rnec7_3286* also harbors three identifiable malonyl-CoA-binding regions.

While several unrelated molecules scored among the top-ranked ligands in docking analyses, acetyl-CoA, acetoacetyl-CoA, HMG-CoA, and other related intermediates did not appear among the lowest G-scores (Table S6). On the other hand, malonyl-CoA and CoA consistently ranked among the top ligands for the primary cavity of both proteins, in agreement with a potential role in chalcone biosynthesis. Chalcones are typically associated with plants, although fungal chalcone biosynthesis has been documented. One of the earliest described fungal T3PKS was a chalcone-flavone isomerase in *Neurospora crassa* (Hashimoto, Nonaka, & Fujii, 2014). More recently, Furumura et al. (2023) described the synthesis of chlorflavonin in *Discosia*, a metabolite also isolated from *Aspergillus* (Wang et al., 2025).

Various other natural chalcone derivatives (4-hydroxychalcone, pinocembrin chalcone, Isoneobavachalcone, Orotinichalcone, and Spinochalcone A) and glycosylated compounds (2′,4,4′,6′-Tetrahydroxychalcone 4′-O-β-D-glucoside, 2′,6′,β-Trihydroxy-4′-methoxychalcone, and 3′-(β-D-Glucopyranosyl)-2′-hydroxy-β-oxodihydrochalcone) were detected in our results, which highlight the association of those compounds with the T3PKS in our results. Other chalcone derivatives are likely synthetic (4′,6′-Dimethoxy-4-dimethylamino-2′-hydroxychalcone, 2′,4′,6′,3,4-Pentahydroxy-3′-geranyl-5-prenyldihydrochalcone, and 2′,4′,6′,β-Tetrahydroxychalcone 2′-glucoside), although little information is available regarding those compounds.

CoA is a central cofactor in numerous biosynthetic pathways, functioning as an acyl group carrier and facilitating the formation of thioester intermediates (Harijan et al., 2023). In chalcone biosynthesis, starter substrates such as p-coumaroyl-CoA require activation via thioester linkage to CoA, enabling iterative condensation with malonyl-CoA catalyzed by chalcone synthases to produce a characteristic 15-carbon polyketide backbone (Futyma et al., 2021). Hence, the identification of CoA among the top docking ligands reinforces the hypothesis that those proteins may catalyze chalcone or stilbene biosynthesis.

Moreover, previous studies have also noted challenges in confidently assigning ligands to putative fungal T3PKS genes in fungi (Kramer, Pimentel-Elardo & Nodwell, 2020; Shi-Kunne et al., 2019). Collectively, our results support the interpretation of *rnec7_3286* and *rnec7_4895* as chalcone synthases while acknowledging that their function as HMG-CoA synthases remains plausible, given its central role in fungal primary metabolism.

## CONCLUSIONS

Xylariaceae is a niche-diverse fungal family characterized by highly variable gene content. Within this family, two T3PKS genes are highly conserved and likely encode chalcone synthases. Although further experimental research is required to identify the compounds synthesized by those proteins, our docking analyses support a role in the biosynthesis of a chalcone-related compound, given the favorable binding of malonyl-CoA, CoA, and related ligands.

In *R. necatrix*, niche specificity appears to be a central factor shaping genome evolution. This broad-spectrum necrotrophic/hemibiotrophic phytopathogen shows extensive gene loss compared to its closest relatives, particularly *Nemania*, which also includes endophytic species. Many of the lost orthologous families are associated with SMGCs, including MFS and ABC transporters. Such losses are likely linked to an altered capacity to export or import bioactive compounds, including processes analogous to drug resistance.

Taken together, our results suggest that the extensive gene losses observed in *R. necatrix* reflect an adaptive strategy to broad-spectrum pathogenicity in plants, distinguishing it from closely related endophytic and saprophytic taxa.

## ACKNOWLEDGEMENTS

This work was supported by Fundação Carlos Chagas Filho de Amparo à Pesquisa do Estado do Rio de Janeiro (FAPERJ), Coordenação de Aperfeiçoamento de Pessoal de Nível Superior - Brasil (CAPES; Finance Code 001), Conselho Nacional de Desenvolvimento Científico e Tecnológico (CNPq) and Programa de Apoio à Pesquisa, Inovação e Cultura (PAPIC - UENF). The funding agencies had no role in the design of the study, collection, analysis, interpretation of data, or writing.

## AUTHOR CONTRIBUTIONS

Gabriel Quintanilha-Peixoto: Data curation, Formal analysis, Writing – original draft, Writing – review & editing. Conceptualization, Funding acquisition, Project administration. Ana Luiza Martins Karl: Data curation, Formal analysis, Writing – original draft, Writing – review & editing. Dayana K. Turquetti-Moraes: Writing – original draft. Felipe Rimes-Casais: Formal analysis, Writing – original draft. Aristóteles Góes-Neto: Writing – original draft, Writing – review & editing. Conceptualization, Funding acquisition. Thiago M. Venancio: Writing – original draft, Writing – review & editing. Conceptualization, Funding acquisition, Project administration, Supervision.

## Supplementary Figures

**Figure S1:**
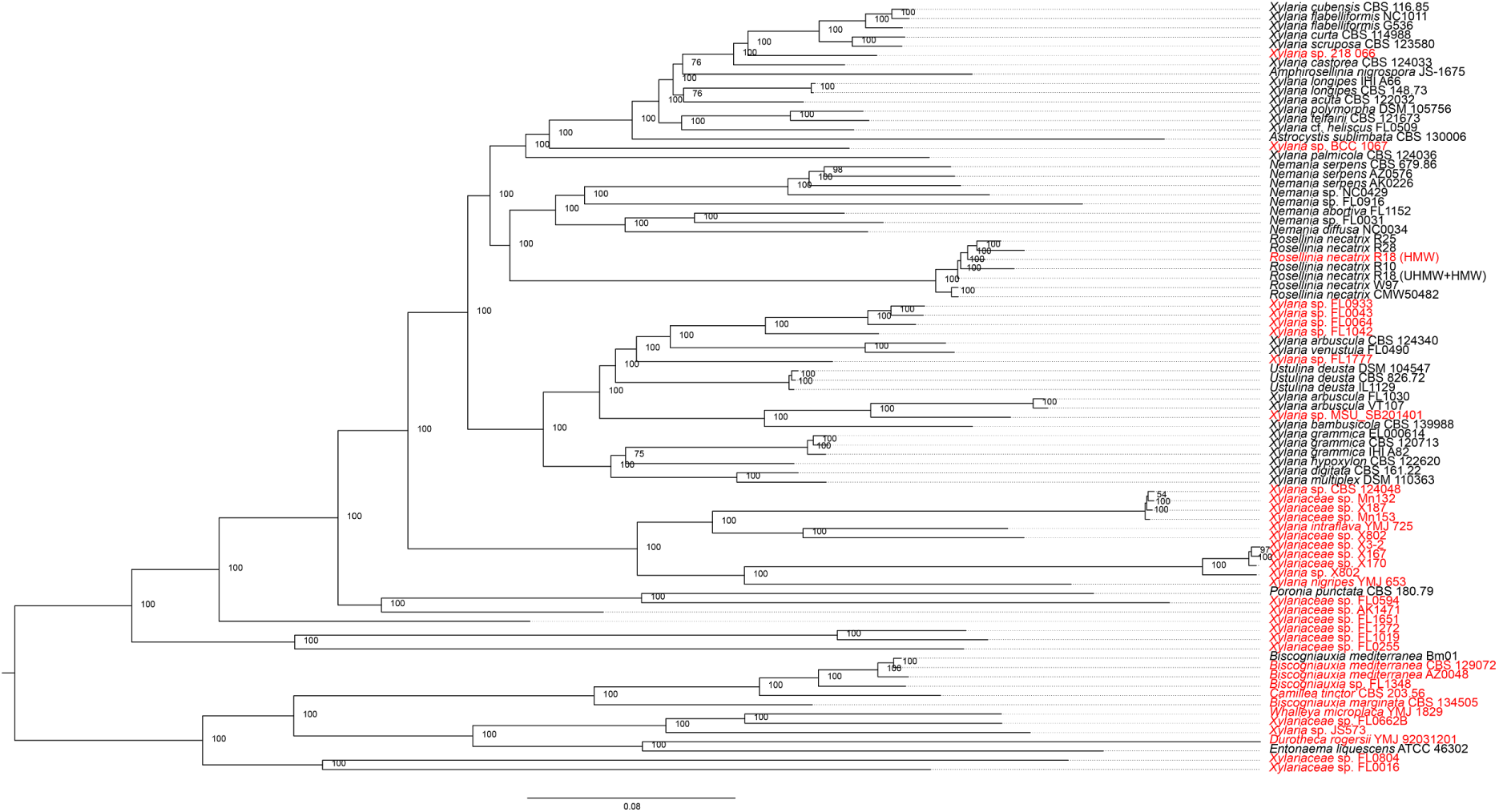
BUSCO phylogenomic tree used for genome filtering.

**Figure S2:**
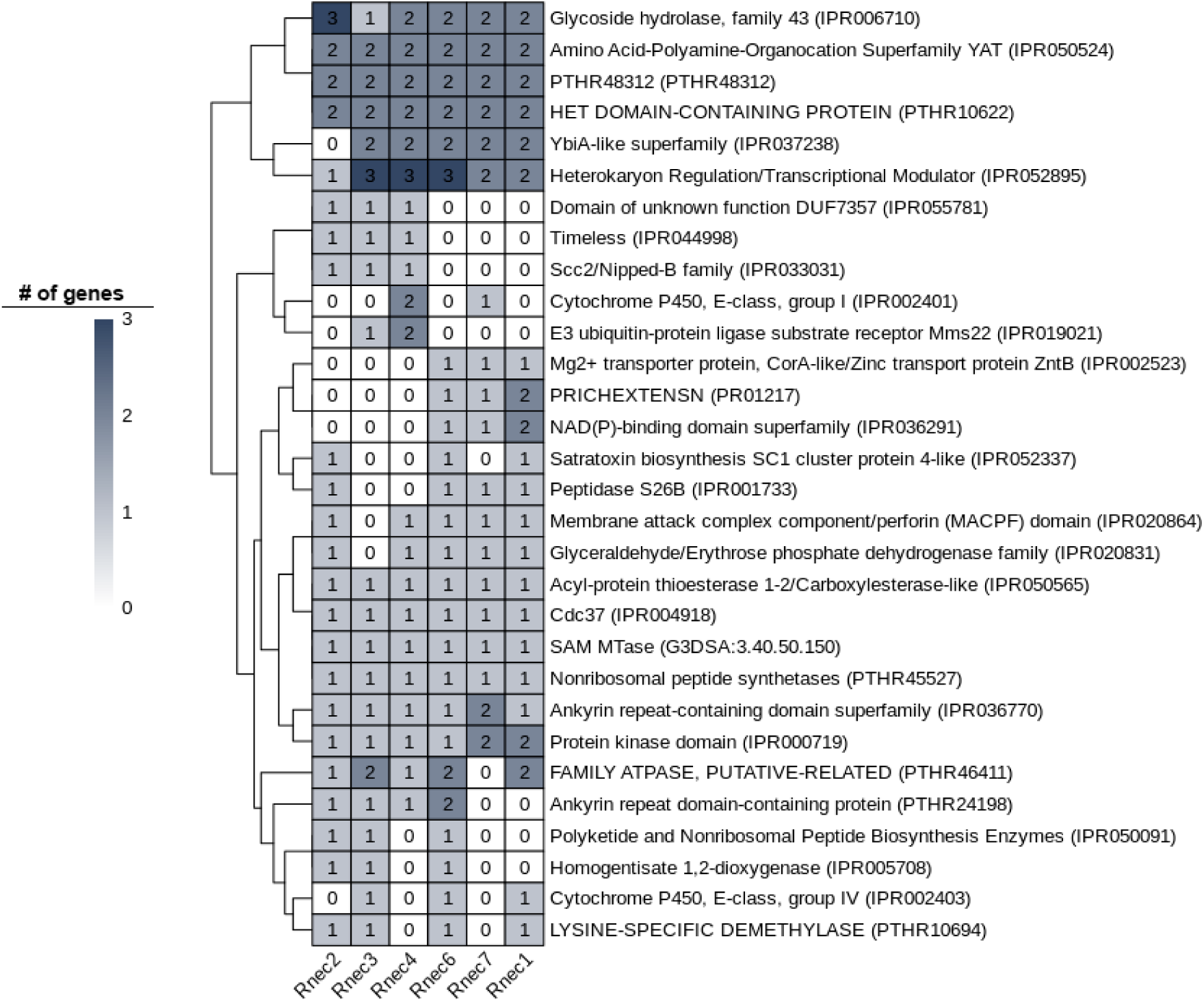
Detailed annotation of the *R. necatrix* cloud genome across different isolates.

**Figure S3:**
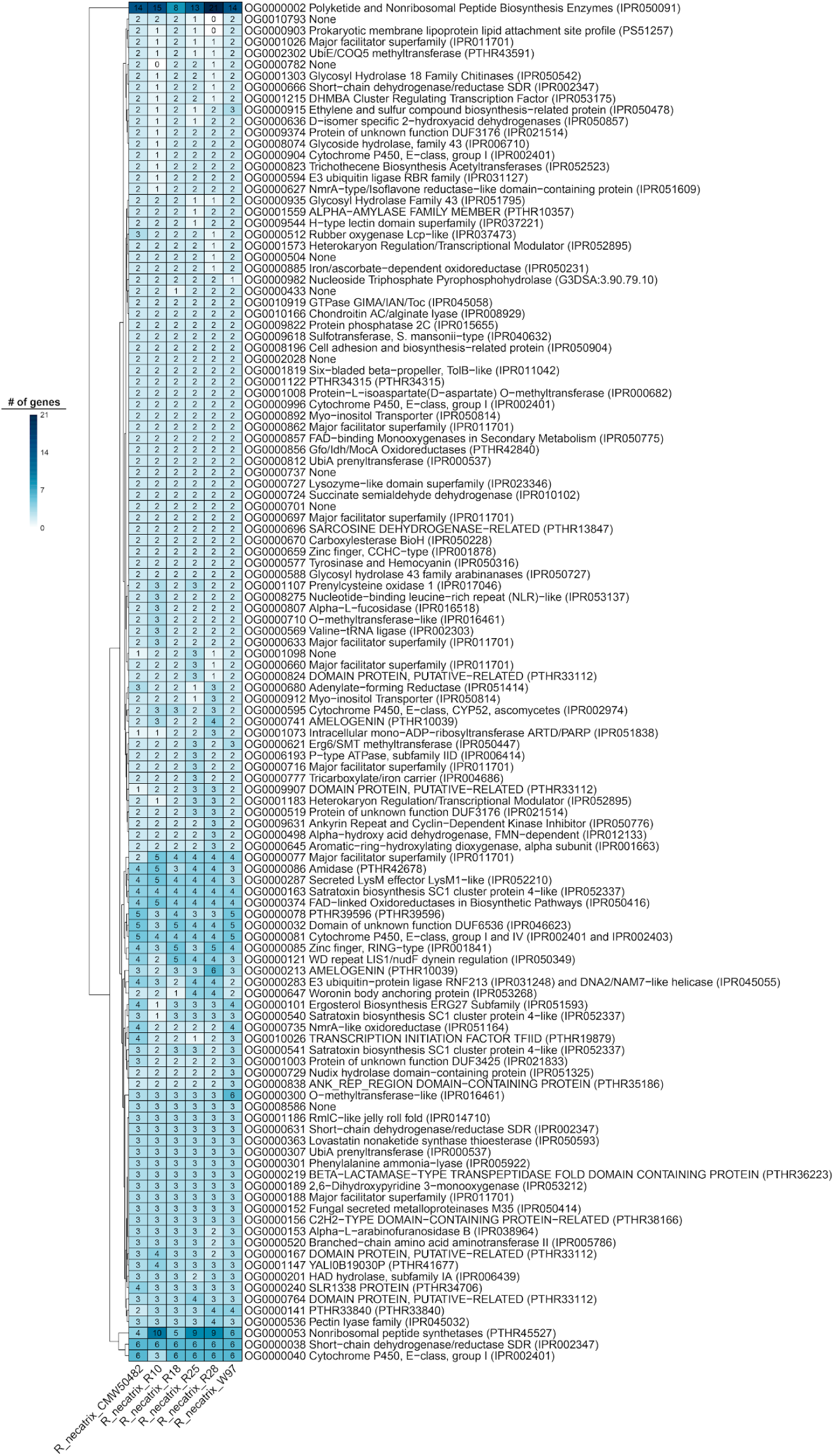
Expanded ortholog gene families and their abundances across *R. necatrix* isolates.

**Figure S4:**
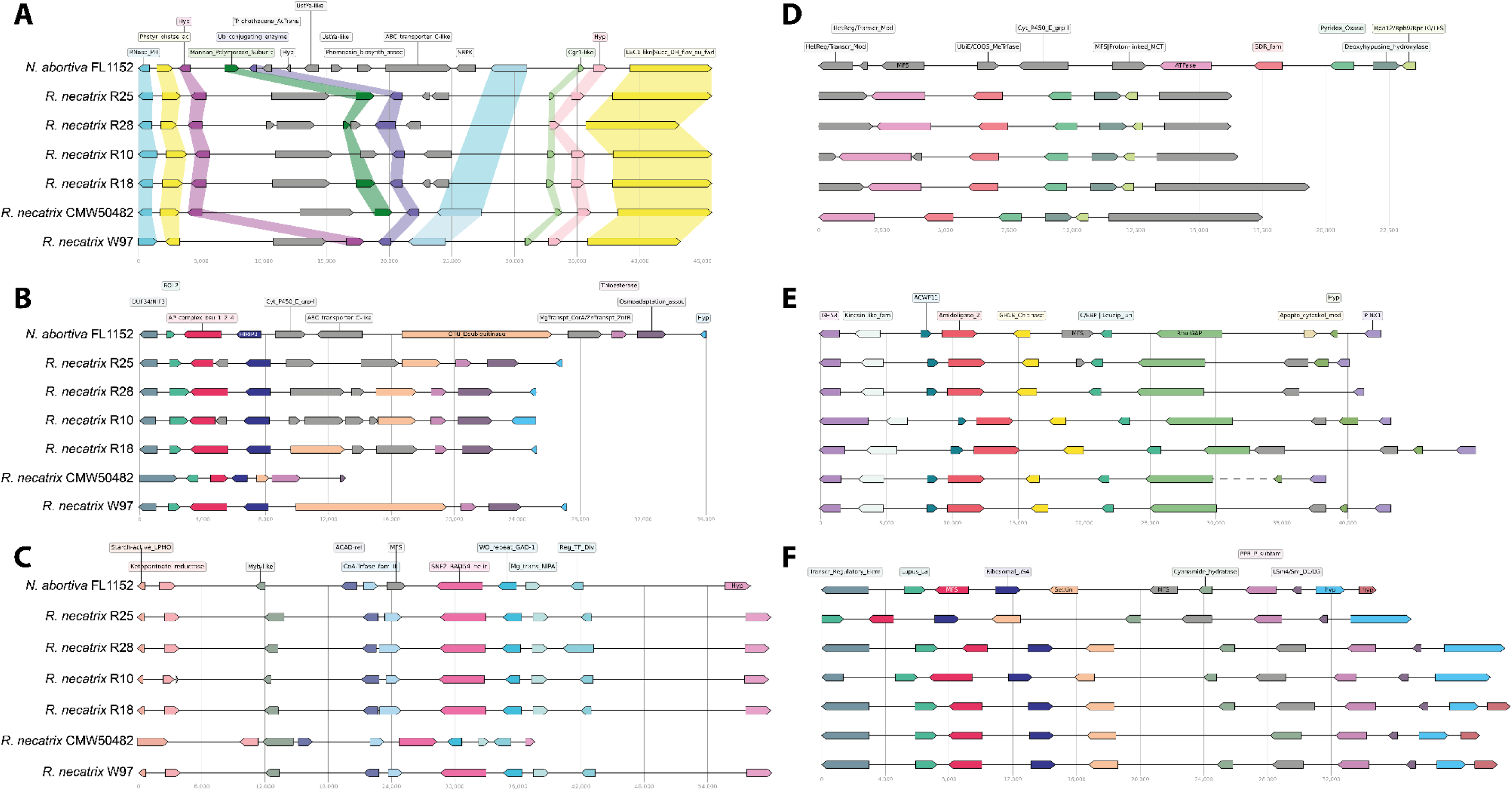
Genomic context around ABC Transporters and MFS gene losses across *R. necatrix* isolates. Similar colors indicate ortholog genes. Grey arrows indicate genes lost in one or more genomes. **A:** OG0000176_3. **B:** OG0000176_4. **C:** OG0000968. **D:** OG0008952. **E:** OG0008993. **F:** OG0009665.

**Figure S5:**
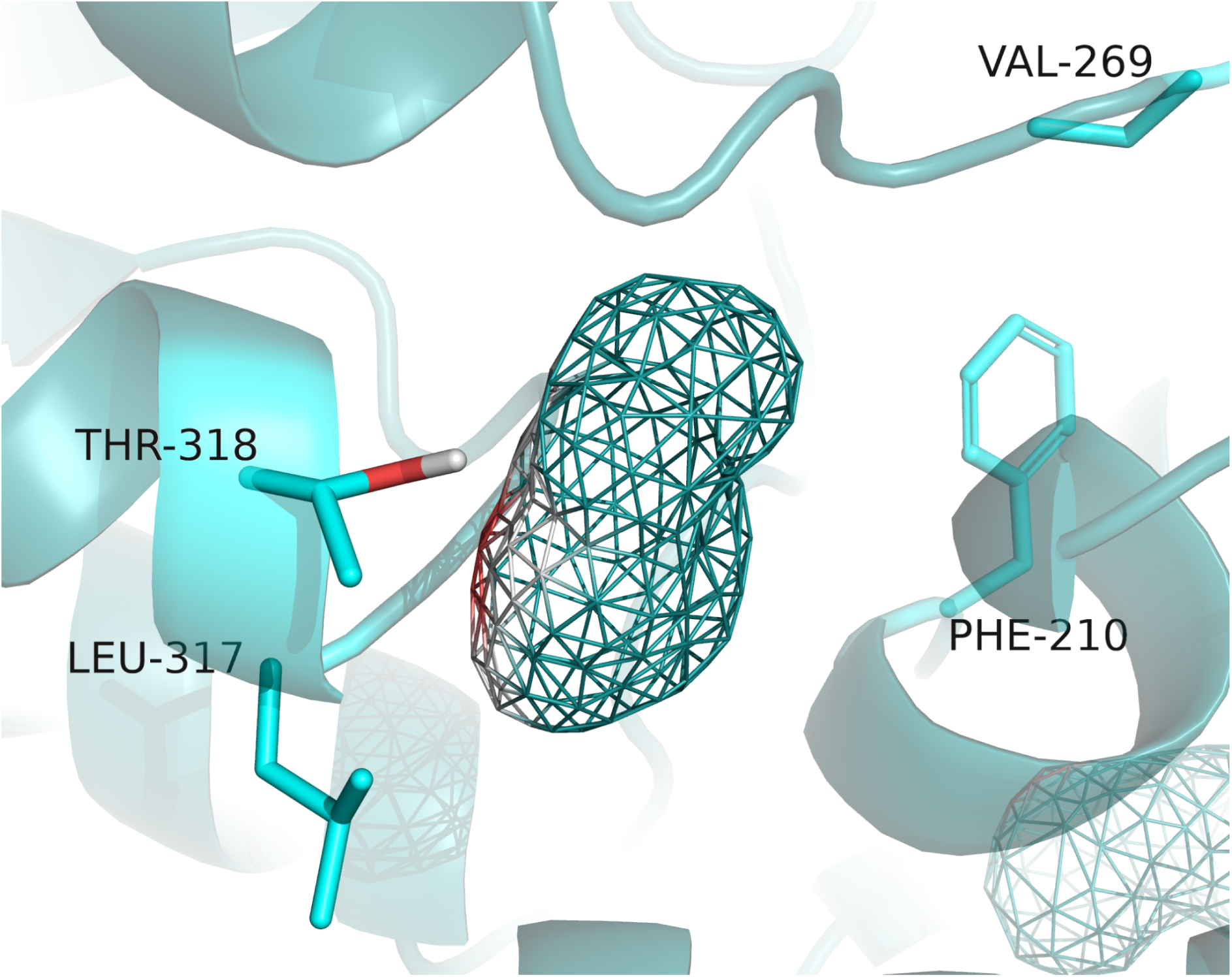
Malonyl-CoA binding site residues initially identified by InterProScan (residues 210, 269, 316-318). Neither CBDock2 nor Maestro recognized this site as a probable docking cavity. Although PyMOL detected it as a cavity, the calculated volume was irrelevant for docking experiments.

